# Model-based concept to extract heart beat-to-beat variations beyond respiratory arrhythmia and baroreflex

**DOI:** 10.1101/2023.03.23.534047

**Authors:** Mindaugas Baranauskas, Rytis Stanikūnas, Eugenijus Kaniušas, Arūnas Lukoševičius

## Abstract

Heart rate (HR) and its variability (HRV) reflect the autonomous nervous system (ANS) modulation, especially sympathovagal balance. This work aims to present concept of a personalizable HR model and an in silico system to identify the HR regulation parameters and subsequently capture residual heart beat-to-beat variations from individual psychophysiological recordings in humans. The model encompasses respiratory sinus arrhythmia (RSA) and baroreflex mechanisms, and uses respiration and blood pressure signals and the time instances of R peaks from an electrocardiogram as inputs. The system extracts the residual displacements of the modeled R peaks relative to the real R peaks. Three components – tonic, spontaneous, and 0.1 Hz changes – can be derived from these R peak residual displacements and can, therefore, enhance HRV analysis beyond RSA and baroreflex. Our model-based concept suggests that these residuals are not merely modeling errors. The proposed method could help to investigate additional neural regulation impulses from the higher-order brain and other influences.

## 1 Introduction

Heart rate (HR) regulation involves several hierarchical levels between the inner nervous system of the heart and the cerebral cortex (Palma and Benarroch, 2014; Schmaußer et al., 2022). Respiratory sinus arrhythmia (RSA) and cardiac baroreflex – the two main lower-order HR regulation mechanisms – can explain the majority of *HR variability (HRV)* (Draghici and Taylor, 2016). However, both overshadow other interesting time-varying influences that can be valuable HRV components. Additionally, respiration rate, age, sex, and other factors must be considered (or controlled) for proper HRV interpretation and comparison between individuals (Laborde et al., 2017; Shaffer and Ginsberg, 2017; Koenig et al., 2021).

Complex underlying regulation mechanisms and personal peculiarities are still not fully disclosed by traditional HRV measures, and other than traditional HRV measures are not always more helpful in the clinical context, nevertheless, research on new ways to evaluate HR dynamics is still encouraged (Sassi et al., 2015). *An alternative approach* could be, for instance, using statistical techniques to invert mathematically formulated forward models (Bach et al., 2018); or fitting HR dynamics to a more comprehensive personalizable HR regulation model to find the weights of regulation mechanisms and analyze the residual HR dynamics unexplained by the model.

Numerous heart regulation models have been developed, for example, by Ursino (1998), its extensions (Ursino and Magosso, 2000; Hsing-Hua Fan and Khoo, 2002; Magosso et al., 2002; Cheng et al., 2010; Albanese et al., 2016; Park et al., 2020), Kotani et al. (2005), its derivations (e.g., Ishbulatov et al., 2020), and other closed-loop models (Van Roon et al., 2004; Ortiz-León et al., 2014) and specific purpose open-loop models (Randall et al., 2019) for humans. Linking models with registered biosignals would make it more adaptive and *personalizable*. However, known models are centered on low-order HR regulation, and this is understandable since regulation at *lower levels* is more deterministic and has been more extensively investigated.

We hypothesize that HR variations unexplained by lower-order mechanisms – neither RSA nor cardiac baroreflex – could contain relatively more influences associated with higher-order brain regulation mechanisms. Activity in some higher brain areas has associations with specific ANS branches, for example, parasympathetic influence could come from the ventromedial prefrontal cortex (Wong et al., 2007; Ziegler et al., 2009; Schmaußer et al., 2022), right (Coote, 2013) or/and anterior (Chouchou et al., 2019) insular cortex, and sympathetic influence could come from the left (Oppenheimer et al., 1992; Coote, 2013) or/and posterior (Chouchou et al., 2019) insular cortex, locus coeruleus (Mather et al., 2017). The effect of a single parasympathetic impulse on HR is swift and could vanish within one second (Spear et al., 1979; Berger et al., 1989), meaning that HR modeling should preferably be done at high temporal resolution to align parasympathetic bursts within the cardiac cycle. However, the known personalization solutions (Olufsen and Ottesen, 2013; Randall et al., 2019) use mean arterial blood pressure (ABP) instead of a continuous ABP signal that changes within the interbeat period. More granular timing could also help extract higher-order HR regulation since brain stimulation correlates with HRV measures sensitive to fast changes in HR, e.g., high-frequency power (Schmaußer et al., 2022), however standard HRV measures usually generalize several minutes of psychophysiological data (Shaffer and Ginsberg, 2017). To our knowledge, no computational psychophysiologically-based personalizable model of HR regulation that considers ANS activity fluctuations within the cardiac cycle has been proposed thus far.

This article *aims* to present concept of a computational model-based method to identify heart rate regulation parameters associated with respiratory arrhythmia and baroreflex to subsequently capture the remaining beat-to-beat variations from individual empirical human psychophysiological recordings.

First, a comprehensive computational HR lower-order neural control model for RSA and baroreflex is constructed. Then, a system for personalization of the model parameters is presented through the use of experimental signals. Results showing residual HR variations beyond RSA and baroreflex, possibly attributable to higher-order brain control, are shown and discussed.

## 2 Methods

The HR regulation model encompasses three main domains: 1) heart, 2) brain lower- and higher-order neural control, and 3) vasculature (see **Figure 1**). The proposed model follows the general scheme of RSA and baroreflexes described by Feher (2017, 613, their Figure 5.13.6) and Jänig (2006, 401). The vagal baroreflex entails baroreceptors, *nucleus tractus solitarius* (NTS), and *nucleus ambiguus* (NAmb); sympathetic baroreflex – baroreceptors, NTS, rostral ventrolateral medulla (RVLM), intermediolateral column (IML). The RSA is mediated mainly by the respiration center, NAmb, X cranial nervus vagus, and sinoatrial node (SAN). Because of its link to the NAmb, the respiratory system can be considered a neural respiration center and thus interpreted as a lower-order neural control. The internal algorithmic structure of most modules of this general scheme is detailed in Section 2.1.

**Figure 1.**
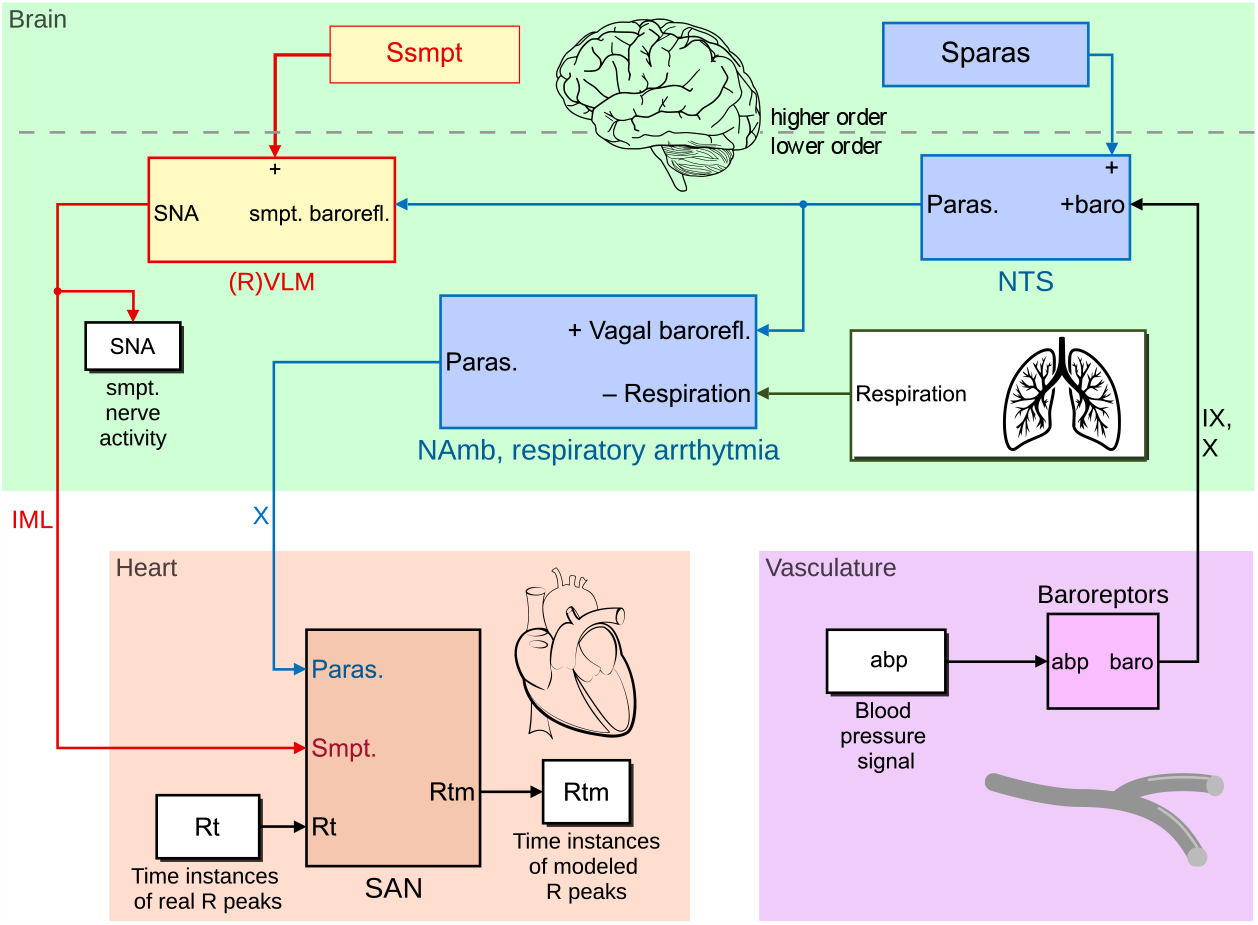
Heart rate regulation model overview scheme in Simulink. It encompasses three main domains: 1) heart (orange), 2) brain (green) – higher-order neural control, also lower-order neural control through respiratory sinus arrhythmia and baroreflex, and 3) vasculature (pink). Abbreviations of structures: IML, intermediolateral column; IX, nervus glossopharyngeus; NAmb, nucleus ambiguus; NTS, nucleus tractus solitarius; (R)VLM, (rostral) ventrolateral medulla; SAN, sinoatrial node; X, nervus vagus. Other abbreviations: baro, activity of baroafferents; paras, parasympathetic activity; smpt, sympathetic activity; Sparas, higher-order parasympathetic activity; Ssmpt, higher-order sympathetic activity. Most modules of this general scheme have a more detailed internal algorithmic structure described in a dedicated subsection.

The proposed HR regulation model uses respiration and ABP signals and the time instances of R peaks from an EKG as inputs to produce modeled R peaks and sympathetic nerve activity (SNA). The model aligns the real and modeled R peaks needed for later extracting the residual HR variations, i.e., the displacements of the modeled R peaks relative to real R peaks. Processed psychophysiological signals (Section 2.3.2) and manually configured set of various parameters (e.g., lower and upper boundaries of the parameters of the HR regulation model), are employed to find the set of model parameters with the lowest error score through optimization algorithms (see **Figure 2** left). The error score is calculated based on the R peak displacements in time and SNA. Section 2.2. describes the system for personalization of the model parameters.

**Figure 2.**
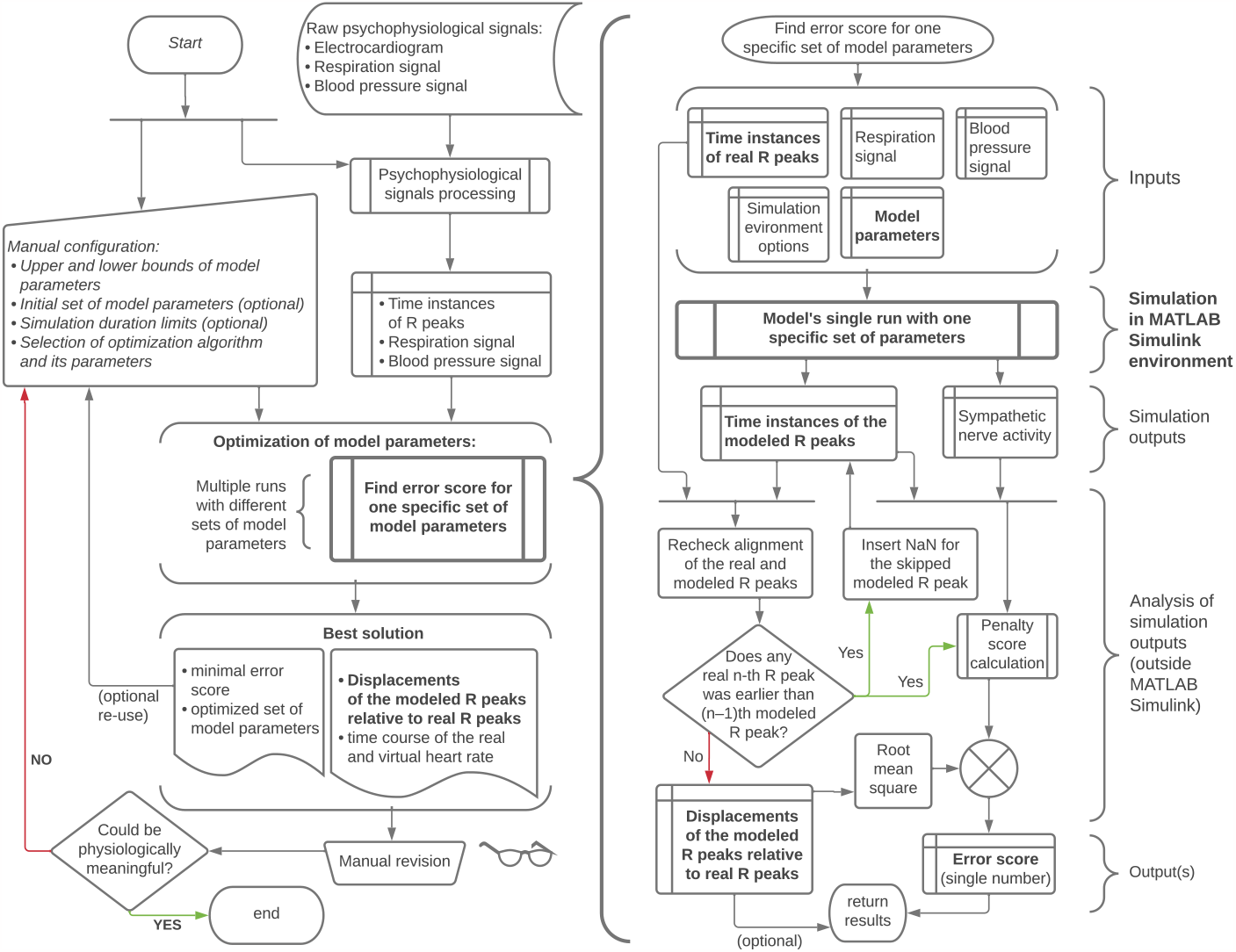
Algorithmic scheme of concept. Here ‘model’ (and “Model’s single run with a specific set of parameters”) refers to the heart rate regulation model realized in MATLAB Simulink environment. Although this model usually returns the modeled R peaks aligned to the real ones, re-check and correction is needed for rare cases of the skipped modeled R peak(s). Abbreviations: NaN, not a number.

The model is implemented in MATLAB Simulink environment and runs with ODE3 (Bogacki-Shampine) solver and fixed step of 1 millisecond. The computational system implementing this model-based concept can be downloaded from the open-source Zenodo repository: <https://doi.org/10.5281/zenodo.7765459>.

### 2.1 Heart rate (HR) regulation model

#### 2.1.1 Heart: Integration of parasympathetic and sympathetic activity in the sinoatrial node

HR would be virtually constant for each individual if it depended only on the SAN by isolating it from external neural impulses, such as by denervation by transplantation or pharmacological blockade (Jose and Collison, 1970; Zeuzem et al., 1991). This *pacemaker* frequency slowly decreases with age (Jose and Collison, 1970). Our HR regulation model incorporates this *pacemaker* frequency in the variable *HRbasal* (see **Figure 3**). If age is known, *HRbasal* can be defined by the Jose and Collison (1970) formula:

**Figure 3.**
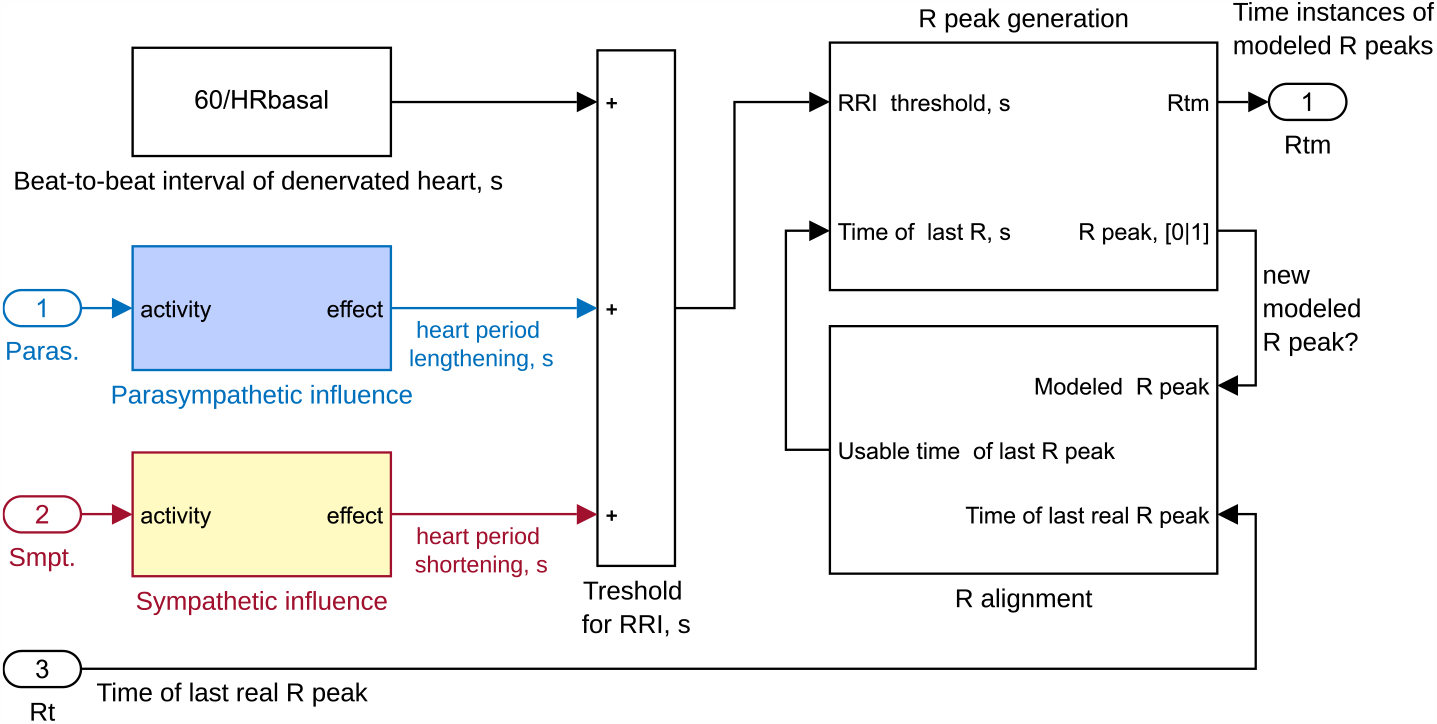
Sinoatrial node block scheme in Simulink. New R peaks are modeled when a time threshold for R-R intervals (RRI) is reached, which is calculated with respect to the real R peak time instances; that is time instances of last real R peak are used to align real and modeled R peaks and find the lack or surplus of threshold for each R peak time during analysis outside the model. Abbreviations: HRbasal – pacemaker’s internal frequency; Paras, parasympathetic activity; Rtm, time instances of modeled R peaks; Smpt, sympathetic activity.

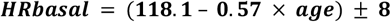

A deviation of up to 8 beats per minute (BPM) of *HRbasal* is allowed for model parameter optimization.

The SAN is innervated by the X cranial nerve (vagus) and IML of the spinal cord that provide *parasympathetic* and *sympathetic input*, respectively (Palma and Benarroch, 2014), as shown in **Figure 1. Figure 3** depicts their placement in our model. Parasympathetic burst decelerates HR with a delay of about 0.2 s, and its most potent effect is at approximately 0.5 s after stimulation, vanishing 1 s afterward. In contrast, a sympathetic burst accelerates the HR with a delay of about 1-2 s (we set 1.5 s), exerting its strongest effect at approximately 4 s and vanishing 20 s after the stimulation (Spear et al., 1979; Berger et al., 1989). We implemented a custom-made transfer function based on Spear (1979, their Figure 2) and Berger et al. (1989, their Figure 4) empirical data for parasympathetic (**Figure 4**) and sympathetic (**Figure 5**) neural burst effect on the heart period. To emulate the gradual effect of the ANS on the heart period before its peak, we use the difference between two transfer functions instead of a single first-order transfer function as employed in the Ursino (1998) and similar models. Single transfer first-order functions suggest an exponential-like ANS effect on HR that is physiologically incorrect. In our SAN block, we use R-R intervals (RRI) instead of HR since the influence of parasympathetic and sympathetic activity (burst frequency) on RRI is almost linear (Draghici and Taylor, 2016; Geus et al., 2019).

**Figure 4.**
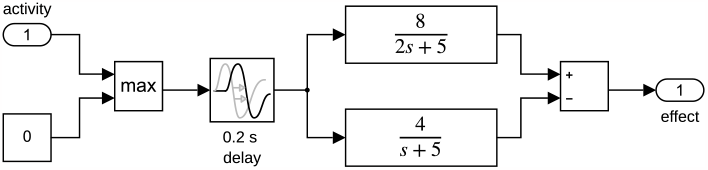
Parasympathetic influence sub-block of sinoatrial node block in Simulink.

**Figure 5.**
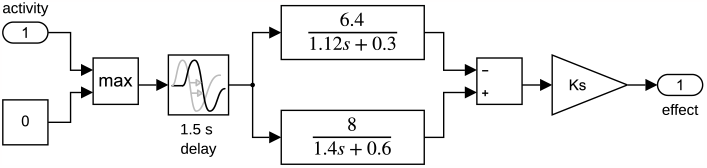
Sympathetic influence sub-block of sinoatrial node block in Simulink. Ks, sympathetic activity coefficient.

The time-varying threshold for RRI is obtained by summing the heart period of the denervated heart (60/*HRbasal*) and the parasympathetic and sympathetic effects mentioned above (see **Figure 3**). When the accumulated time after the last R peak is more than 300 ms and hits the dynamically changing RRI threshold, the pacemaker fires (a new R peak is generated) and is immediately reset to zero in the “EKG R peak generation” sub-block (see **Figure 3** and **Figure 6**). Here, time instances of modeled R peaks are stored in memory for analysis outside the model.

The crucial part of this HR model is the sub-block for aligning the real and modeled R peaks (see **Figure 6** for its context within the SAN sub-block and **Figure 7** for its detailed implementation in Simulink). Usually, this block returns the time instance of the previous real R peak as output for the corresponding modeled R peak. However, if the modeled R peak precedes the real R peak, then infinity will be temporarily returned as the “usable time of the last R peak”, i.e. the new modeled R peaks are then blocked until reset by the following real R peak. If the real R peak precedes the modeled one, then the update of the output value will be temporarily blocked until a new modeled R peak. This allows modeling R peaks relative to the real R peaks. In addition, this approach enables the extraction of the R displacements by comparing the real and the modeled R peaks outside the model. However, this sub-block cannot deal with cases when the modeled R peak happens later than one real R peak; the additional re-check and correction of these cases outside the model is described in Section 2.2.

**Figure 6.**
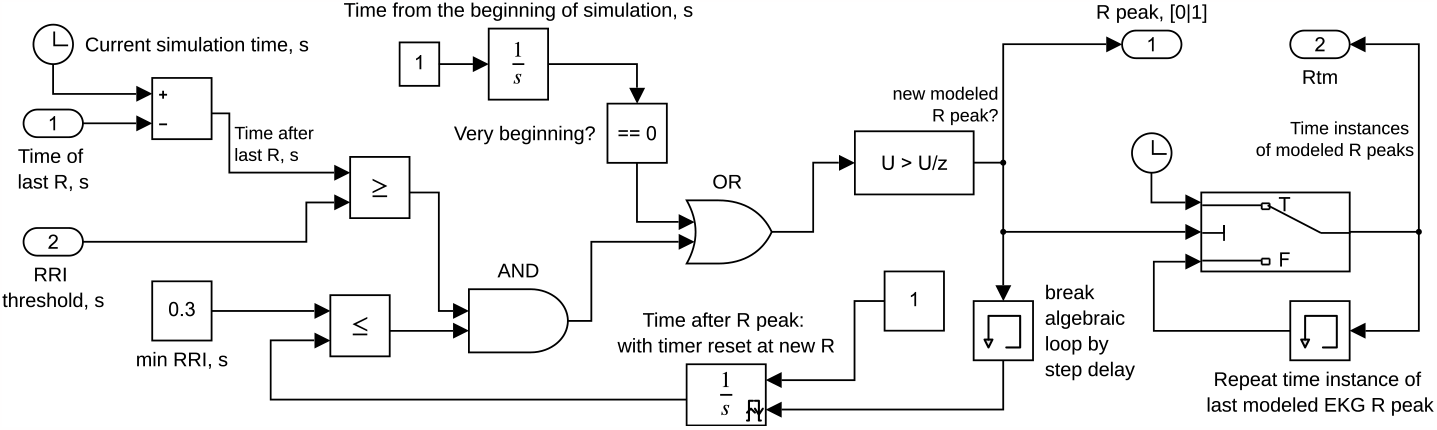
EKG R peak generation sub-block in Simulink. When the accumulated time after the last R peak is more than 300 ms and hits the dynamically changing R-R interval (RRI) threshold, the pacemaker fires (a new R peak is generated) and is immediately reset to zero. Note 1: ‘current simulation time’ and ‘time from the beginning of simulation’ will not be the same if the simulation starts not at zero seconds of psychophysiological recording. Note 2: ‘time instances of modeled R peaks’ are not the same as ‘time of last R’ because R alignment is enabled outside this sub-block. Time instances of modeled R peaks are passed to be stored in memory (here ‘Rtm’) for analysis outside the model.

**Figure 7.**
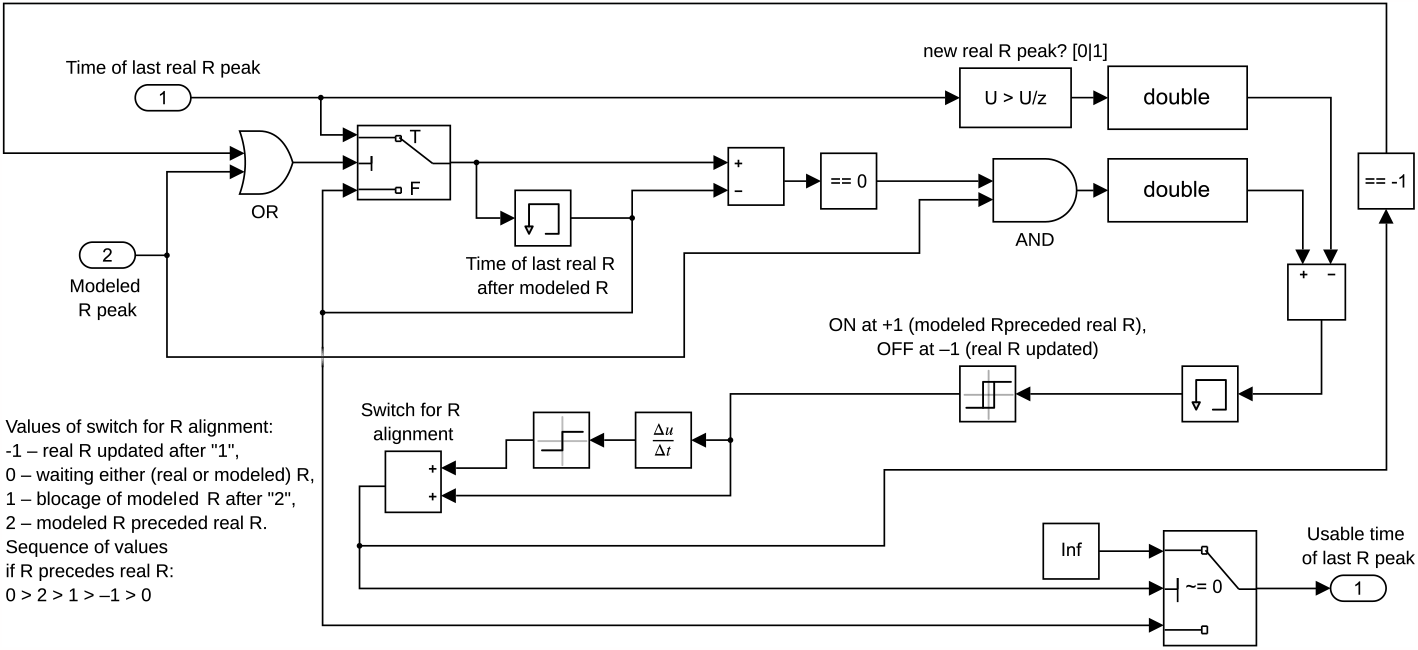
Simulink sub-block for alignment of the real and modeled R peaks.

#### 2.1.2 Vasculature: Baroreception

The blood pulse wave stretches the vessel walls. It reaches the arterial baroreceptors in the aortic arch after 10–15 ms and the carotid sinus after 40–65 ms, corresponding to about 90 ms and 140 ms after the R peak in the EKG, respectively (Edwards et al., 2009; Wehrwein and Joyner, 2013; Martins et al., 2014). Although stimulation of aortic and carotid baroreceptors can have different impacts on the HR control (Ishii et al., 2015), we do not differentiate them. We modeled the integrated baroreceptor activity and set an average delay of 30 ms (see **Figure 8**). Some baroreceptors react more to phasic changes of ABP, while others respond more to the absolute level of ABP. Thus, for integration, we add the derivative of ABP by the coefficient *Pk* and scale the resulting sum by *Kb*. Baroreceptors may take about 20 ms to react to ABP (Borst and Karemaker, 1983), and the integrated baroreceptor activity peak is estimated at about 250 ms and extinguishes until 390 ms after the EKG R peak (Edwards et al., 2009). Thus, to match this with experimental or simulated data, a 50 ms time lag is added for modified ABP signal *P* in our model. Regardless of their place, baroreceptors have different activation thresholds, response functions, and saturation levels Field (Wehrwein and Joyner, 2013; Feher, 2017, 610). To mimic the varying baroafferent activity (*baro*) within the cardiac cycle, we use a sigmoidal function similar to the one in the Ursino (1998) model:

**Figure 8.**
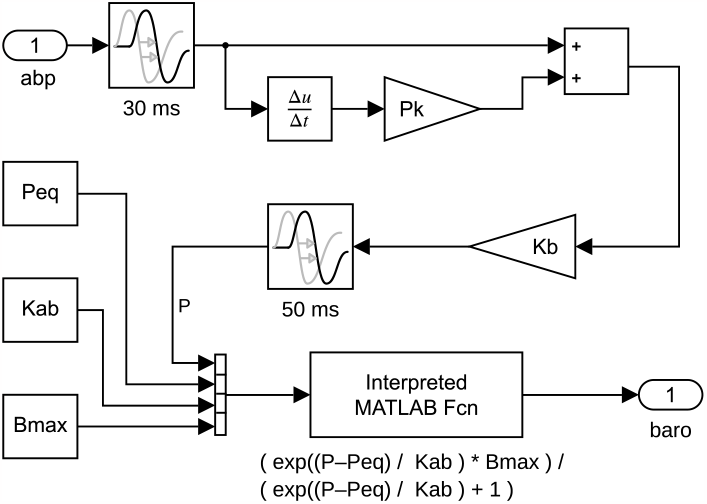
Simulation of the integrated baroreceptor responses (baro) from arterial blood pressure (abp) signal in Simulink. Abbreviations: Bmax, saturation level of baroafferent activity; Kab, the slope-related coefficient for baroreception; Kb, scaling factor of *abp* and its derivative sum for baroreception; Peq, the central point of the sigmoidal functional for baroreception; Pk, *abp* derivative coefficient for baroreception.

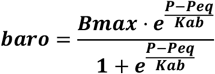

where *Peq* is the central point of the sigmoidal functional, *Kab* is the slope-related coefficient, and *Bmax* is the saturation level, which can be sensitive to postural position. Unlike Ursino (1998), we did not set a minimum activation level since most baroreceptors do not show spontaneous activity. Instead, we set the tonic parasympathetic activity from the higher-order brain areas to NTS (see **Figure 1**). The values of baroreception parameters are suggested to be selected so that the baroreceptors would be more sensitive to the main pulse front and less sensitive to the dicrotic notch.

#### 2.1.3 Brain

##### 2.1.3.1 The path through NTS for cardiac vagal and sympathetic baroreflexes

Baroreceptive information from the carotid and aortic baroreceptors travels via the cranial nerves (glossopharyngeal and vagus, respectively), reaching the NTS within 10–15 ms (Wehrwein and Joyner, 2013; Martins et al., 2014). NTS is the first central relay station of visceral afferent information and is part of the circuit for all medullary reflexes, including the baroreflex (Palma and Benarroch, 2014). NTS projects within 100 to 150 ms to areas responsible for baroreflex effects, including NAmb and RVLM, which is about 200 to 300 ms after the R peak in the EKG (Edwards et al., 2001). Thus, we add 15 and 125 ms time delays, respectively, as the fixed values in the NTS scheme (see **Figure 9**).

**Figure 9.**
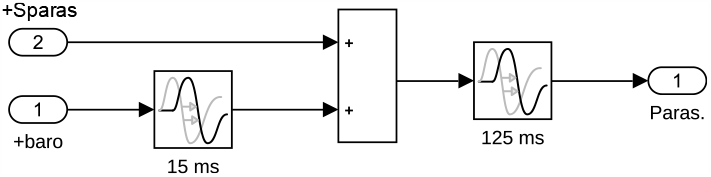
Nucleus tractus solitarius scheme in Simulink. Abbreviations: baro, activity of baroafferents; Paras, parasympathetic activity for output; Sparas, higher-order parasympathetic activity for input.

NTS is under the powerful control of the brain and integrates afferent information, sending projections back almost to every great variety of neural structures, including the brainstem, amygdala, and (via thalamus) insular and prefrontal cortices (Jänig, 2006, 311–317). In this HR model, the NTS is set not to have spontaneous activity and to receive the integrated parasympathetic excitatory input *Sparas* from the “brain” (see **Figure 1** and **Figure 9**). NTS is also the only structure to receive this input.

Baroreflex acts via NTS in three ways: as cardiac vagal (parasympathetic) baroreflex (via NAmb, see 2.1.3.2 subsection), cardiac sympathetic baroreflex (via RVLM, see 2.1.3.3 subsection), and noncardiac sympathetic baroreflex (not relevant to our HR model). In the cardiac vagal baroreflex circuit, NTS directly excites preganglionic cardiac vagal neurons in NAmb, resulting in HR deceleration (Wehrwein and Joyner, 2013; Palma and Benarroch, 2014).

##### 2.1.3.2 Preganglionic cardiac vagal neurons in the nucleus ambiguus (NAmb) and respiratory sinus arrhythmia (RSA)

Most (about 80%) of preganglionic cardiac neurons originate from NAmb and can effectively decelerate HR via myelinated fibers, whose activity is usually vastly similar to the respiration pattern (Coote, 2013). The effect of other cardiac vagal neurons on the HR is relatively small, more delayed, and slower than those of NAmb (Coote, 2013). NAmb neurons are not spontaneously active themselves. They are excited by other areas of the brainstem – predominantly by the NTS – and are inhibited by neurons of the medullary ventral respiratory group (“respiration center”) only during inspiration, i.e., they are not blocked during expiration or holding of breath, apnea (Dergacheva et al., 2010; Palma and Benarroch, 2014; Feher, 2017, 612–613). Nevertheless, this inhibition is not total and does not imply the absence of inhibition later (Geus et al., 2019). The aggregated effect causes RSA – a temporal withdrawal of the predominant parasympathetic activity that accelerates HR during inspiration. Therefore, we include only preganglionic cardiac neurons of NAmb with excitatory input from NTS (incorporating baroreceptive information) and inhibition by multiplication of the square root of the derivative of the respiration signal and transfer function with *Krsa* and *Drsa* parameters only at inspiration (see **Figure 10**). The NAmb output activity is set to be always non-negative.

**Figure 10.**
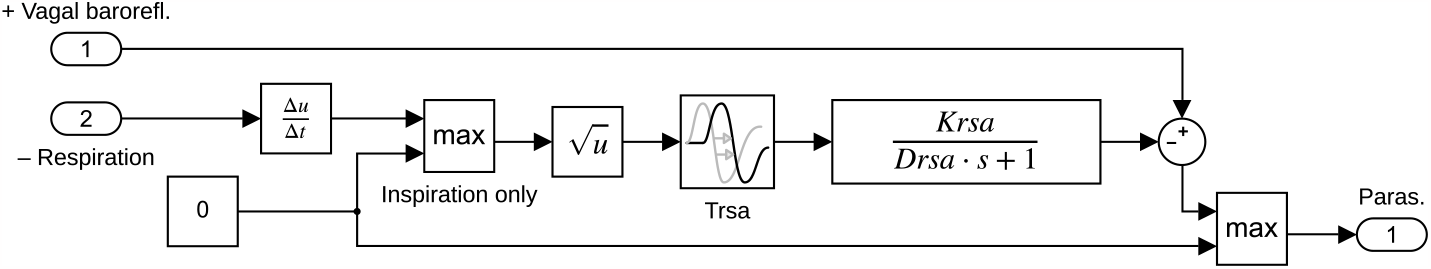
Nucleus ambiguus scheme for *respiratory arrhythmia* in Simulink. Parasympathetic activity (‘Vagal baroreflex’ input #1) is inhibited during inspiration with little time delay *Trsa*. Inhibition strength depends on respiration (input #2) signal derivative (its square root), transfer function (where *Drsa* is in the denominator), and custom *Krsa* coefficient. Block outputs parasympathetic activity (Paras.).

##### 2.1.3.3 Rostral ventrolateral medulla (RVLM) for sympathetic baroreflex

The aggregated effect of baroreceptor stimulation inhibits sympathetic activity. In the sympathetic baroreflex circuit, NTS through the caudal ventrolateral medulla (CVLM) or directly inhibits RVLM (Wehrwein and Joyner, 2013; Palma and Benarroch, 2014). Thus, in our model overview scheme, NTS is shown to act on “(R)VLM” (see **Figure 1**).

Sympathetic baroreflex does not produce sympathetic outflow by itself: if the central sympathetic activity is low or insufficient (e.g., during anesthesia), sympathetic baroreflex will not be noticeable (Karemaker, 2022). RVLM has spontaneous activity and receives various excitatory inputs (Jänig, 2006, 393–396). Sympathoexcitatory RVLM neurons are activated by psychological stress, pain, hypoxia, hypovolemia, and hypoglycemia both directly and via descending inputs from higher-order structures (e.g., insula, prefrontal cortex, amygdala, and hypothalamic subnuclei) (Palma and Benarroch, 2014; Barman and Yates, 2017). In this HR model, the RVLM theoretically has constant spontaneous activity *Arvlm_sp* and receives the integrated sympathetic excitatory input from “brain” *Ssmpt* (see **Figure 11** left, and subsection 2.1.3.4); RVLM is also the only structure receiving such input.

**Figure 11.**
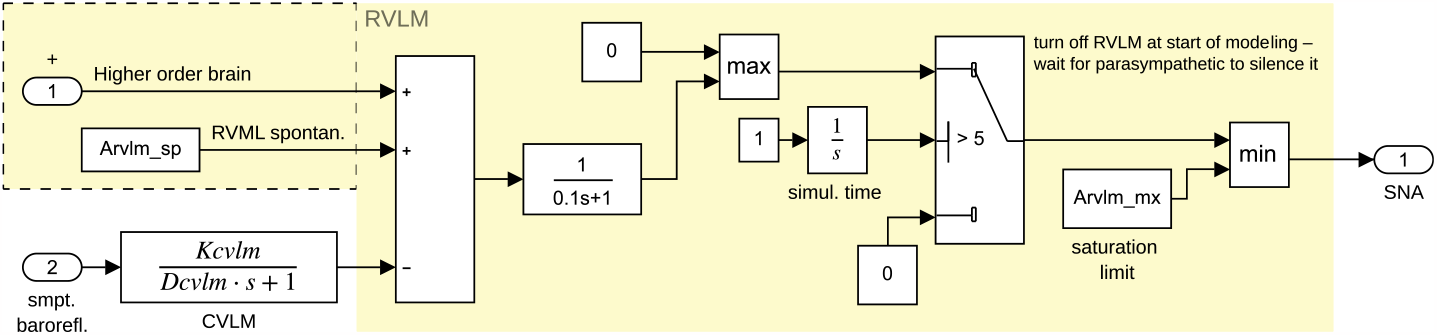
Rostral ventrolateral medulla (RVLM) algorithmic structure in Simulink to generate pulsatile sympathetic nerve activity (SNA) when it is not fully blocked by the caudal ventrolateral medulla (CVLM). Variables: *Arvlm_sp* – constant spontaneous RVLM internal activity, *Arvlm_mx* – saturation limit of RVML output; *Dcvml* and *Kcvml* – NTS parasympathetic effect on sympathetic activity coefficient.

The RVLM output has a pulsatile pattern that becomes visible when NTS/CVLM does not block it. The NTS/CVLM inhibitory activity is short, so an additional mechanism of inhibitory inertness seems to exist to keep RVLM pulses short. Since the mechanism by which this happens in the ventrolateral medulla is not yet understood, we implement two first-order transfer functions to obtain a comparable effect. Only one of these transfer functions has variable parameters (*Kcvlm* and *Dcvlm*) and is named “CVLM” (see **Figure 11**).

The RVLM is temporarily turned off for 5 seconds until parasympathetic activity through NTS/CVLM would regularly inhibit RVLM and limit RVLM output to saturation value *Arvlm_mx* (see **Figure 11**, right). If not turned off, the model would produce long-lasting, strong sympathetic effects on HR and vasculature in the beginning of simulation.

The RVLM-generated SNA travels through the sympathetic IML of the spinal cord. IML neurons serve as a common sympathetic effector and through sympathetic ganglia innervate the heart (Wehrwein and Joyner, 2013; Palma and Benarroch, 2014). IML also innervates blood vessels, however arterial (non-cardiac) sympathetic baroreflex is irrelevant to our HR model since the recorded ABP signal is used instead of simulated. Section 2.1.1 describes the cardiac sympathetic baroreflex effect on the withdrawal of sympathetic HR acceleration.

##### 2.1.3.4 Higher-order regulation

The intrinsic HR is about 40 BPM faster than the mean HR in the resting state due to the dominance of the parasympathetic influence (Smetana and Malik, 2013). Higher-order brain regulation is critical to maintaining it, e.g., by salience network (where anterior insular and anterior cingulate cortices are the core of it) (Guo et al., 2016) and (pre-)frontal cortex (Wong et al., 2007; Thayer and Lane, 2009; Ziegler et al., 2009; Smith et al., 2017). The mere thickness of the prefrontal cortex and insula is positively correlated with the resting HRV (Koenig et al., 2021). This higher-order parasympathetic tonic activity is included in our model as the *Sparas* parameter (see **Figure 1**).

Theoretically, the sympathetic tonic activity could be set as the *Ssmpt* parameter (see **Figure 1**). However, during parameter optimization, it is not possible to differentiate between *Arvlm_sp* and *Ssmpt*; thus, we practically set *Ssmpt* = 0, and *Arvlm_sp* means a constant sum of RVLM spontaneous activity and sympathetic excitatory input from the “brain” (see **Figure 11** and sub-chapter 2.1.3.3 above).

### 2.2 System for model personalization

Although the HR regulation model usually returns the correctly aligned real and modeled R peaks, rechecking and correction are needed for the rare cases of skipped modeled R peak: if the real n^th^ R peak was earlier than the (n–1)^th^ modeled R peak, then *NaN (not a number)* must be inserted for the skipped modeled R peak (see **Figure 2**, right).

Minimization of the error score is used for optimizing the parameters. The error score is based on the displacements of the modeled R peaks relative to the real R peaks and the SNA. It is calculated only for a period starting 30 seconds after the beginning of the simulation. The error score is the sum of two components listed below (see **Figure 2**, right):

1. *Root mean square (RMS) of R peaks displacements*, i.e., of the time instance differences (in milliseconds) between the real and the corresponding aligned modeled R peaks;
2. *Penalty score* that consists of:
  - A penalty of 200 × the ratio of the skipped modeled R peaks and all real R peaks within the analysis period; this penalty is added only if this ratio > 5%;
  - A penalty consisting of a sum of penalties (up to a maximum of 250) of the following integrated SNA properties:
    - A penalty of 50 if any SNA lower peak is > 0;
    - A penalty of 50 if SNA saturates for longer than 0.3 s uninterruptedly;
    - A penalty of 50 if the ratio of unique SNA values and the number of all real R peaks within the analysis period < 10%;
    - A penalty of 100 × (time between SNA peaks and the R peaks as found by cross-correlation minus 0.8) – only if the SNA peaks are greater than 0.8 s after the R peaks;
    - A penalty of 200 if SNA is zero all the time.

Several invocations of the optimization algorithm may be required to find the most acceptable local minimum (see **Figure 2**, left). This is because optimization algorithms seldom find the global minimum, and the lowest local minimum may not be psychophysiologically meaningful due to either the morphology of the modeled R peaks or the combination of parameters (e.g., the algorithm gets “stuck” in the lower or upper boundaries). Therefore, the best-optimized solutions must be manually reviewed.

To personalize the HR model parameters, we selected the initial bounds of parameters as disclosed in **Table 1**. A total of 15 (-16) model parameters can be personalized (the 16^th^ *Ssmpt* parameter is theoretical and practically not used). Lower and upper boundaries or initial values for several of them could be roughly estimated by physiological norms (e.g., *HRbasal* through age) or from other models with similar components (e.g., *Peq* and *Kab* from Ursino’s (1998) model). The boundaries for the remaining parameters are set according to our internal manual research (see **Table 1**). These boundaries were defined to prevent the algorithm from getting “stuck” during parameter optimization and limit its search range. Broader search ranges often lead to the wrong optimization result, e.g., the modeled HR being constant equal to the mean real HR.

**Table 1.**
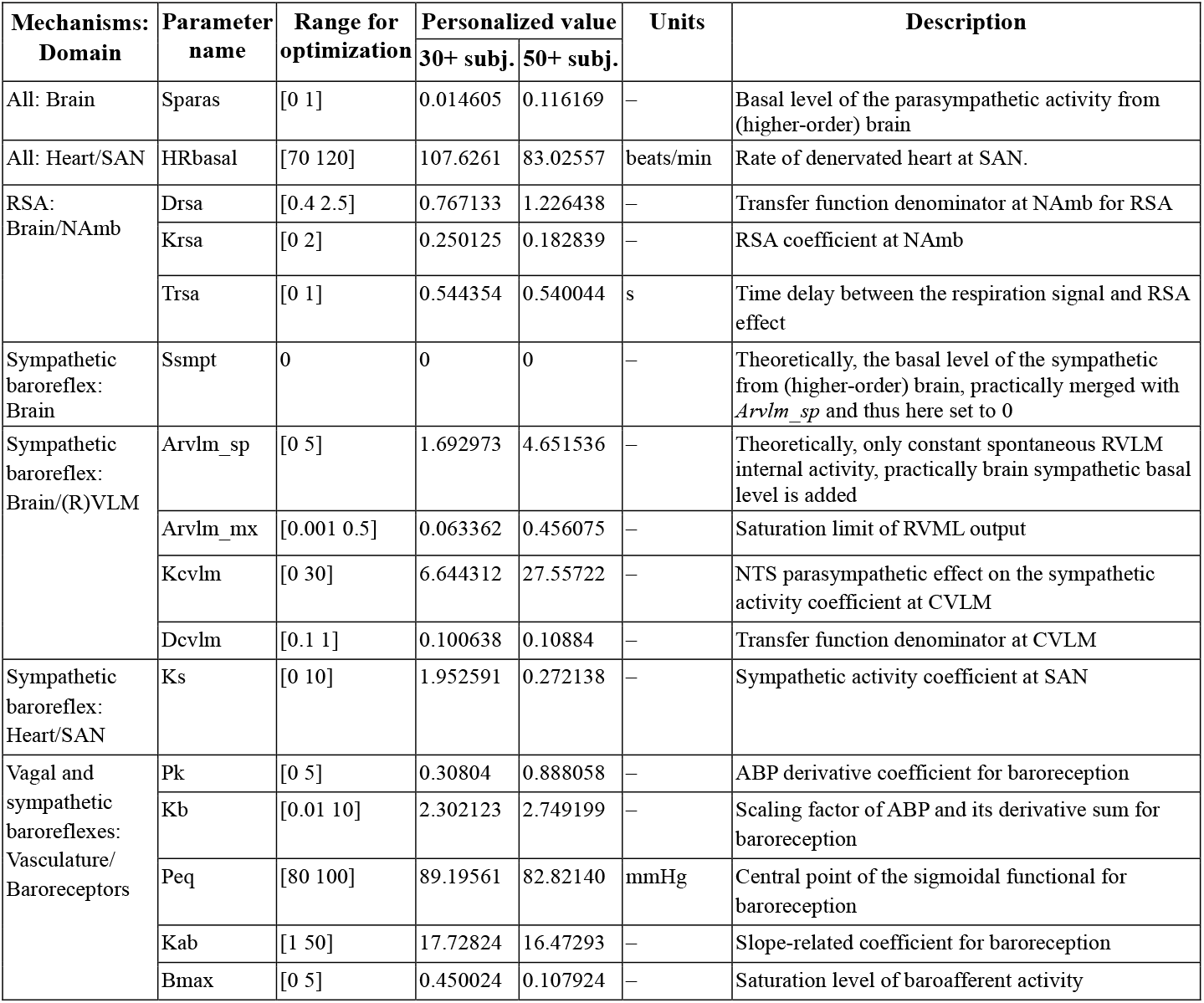
Summary of the parameters of the personalizable heart rate regulation model. Abbreviations: ABP, arterial (blood) pressure; CVLM, caudal ventrolateral medulla; NAmb, nucleus ambiguus; NTS, nucleus tractus solitarius; RSA, respiratory sinus arrhythmia; (R)VLM, (rostral) ventrolateral medulla; SAN, sinoatrial node. Subjects: ‘30+ subj.’ is a tricenarian, ‘50+ subj.’ is a quinquagenarian.

The parameters of the HR regulation model are optimized with MATLAB *particleswarm* and *patternsearch* algorithms from Global Optimization Toolbox. First, for the initial random selection of model parameters and later, for finding the more precise solution with the lowest error score. The default values of ‘*patternsearch*’ optimization algorithm *parameters* must be changed to ensure a successful optimization: (1) the function tolerance (‘FunctionTolerance’) is set to 0.001 to stop iterations once the change in function value reaches less than 0.001; (2) the tolerance on the independent variable (‘StepTolerance’) is set to 0.001 to stop iterations once the distance between two consecutive points is less than 0.001 and the mesh size is less than 0.001; (3) the polling strategy used in the pattern search (‘PollMethod’) is set to GSSPositiveBasis2N, which is more efficient than the default GPSPositiveBasis2N when the current point is near a linear constraint boundary; (4) the order of poll directions in pattern search (‘PollOrderAlgorithm’) changed to “*Random*”; (5) complete poll around the current point (‘UseCompletePoll’). The parameter of the ‘*particleswarm*’ optimization algorithm changed from the default: ‘FunctionTolerance’ set to 0.001, i.e., to end iterations when the relative change in the best objective function value over the last 20 iterations is less than 0.001. One invocation of the selected optimization algorithm runs the HR regulation model multiple times, each time with a different set of HR model parameters.

### 2.3 Testing of a model-based concept

#### 2.3.1 Participants, equipment, and procedure

Two healthy male participants (a tricenarian and a quinquagenarian, without heart-related problems) volunteered for the pilot recordings to fit our HR model. These pilot recordings were conducted in accordance with the ethical principles of the Declaration of Helsinki, including written informed consent and voluntary participation.

Psychophysiological recordings were performed with BIOPAC MP150 (Biopac Systems Inc., 42 Aero Camino, Goleta, CA, United States) with ECG electrodes placed according to lead II and a respiration belt on the abdomen. Using finger cuffs, ABP was registered with the Portapres device (Finapres Medical Systems BV, Palatijn 3, Enschede, Netherlands).

The participants laid in the supine position for 10 min of rest, followed by 3 min of slow breathing, and again 10 min of rest.

#### 2.3.2 Offline signal processing

The *EKG* signal was filtered offline using a bandpass (0.4–35 Hz) Butterworth filter. The time instances of EKG R peaks were identified with a modified Pan-Tompkins (1985) algorithm and then visually inspected. Wrongly detected R peaks were manually corrected by selecting the real EKG R time as signal peak or interpolated.

*The respiratory* signal was filtered with a low-pass filter with a cut-off frequency of 0.4 Hz and normalized between –0.5 and 0.5.

Since *ABP was* registered on the hand fingers, the ABP signal was shifted by –100 ms to compensate for the blood arrival time difference between baroreceptors and fingers. The ABP signal was filtered using a low-pass filter with a cut-off of 10 Hz. The baseline was changed to 80 mmHg.

#### 2.3.3 Derived psychophysiological data

Additional psychophysiological data were derived for more comprehensive visualization of the results. The real HR was directly derived from time instances of the R peaks to see the HR variations along with the respiration signal. Meanwhile, virtual HR was derived from the difference between the real RRI and the corresponding R peak displacements instead of using RRI from the modeled R peaks. To see the associations of ABP with baroreflex-mediated SNA and HR variations, systolic and diastolic blood pressures were derived as upper and lower envelopes of the processed ABP signal. The standard deviation of RRI (SDNN) for the entire 23 min record was computed to evaluate general HRV.

#### 2.3.4 Additional details on psychophysiological data fitting to the model

For parameter personalization, the HR model has been run across the entire 23-minute length of empirical psychophysiological records on MATLAB R2022b parallel pool of 45 processes in a computer with AMD Ryzen Threadripper PRO 5995WX 64-Cores 2.70 GHz processor, 128 GB RAM.

## 3 Results

The processed experimental psychophysiological data were fitted to the HR regulation model by evaluating multiple sets of parameters to find a solution with minimal error score, i.e., the sum of minimal RMS of R peak displacements and the penalty score.

The *particleswarm* algorithm was called twice, and the *patternsearch* algorithm once for optimization of the HR model parameters, passing the best-yielded solution after each invocation to the subsequent optimization as the initial set. For the parameter personalization of a tricenarian, 11859 evaluations of sets of parameters were computed in Simulink for about 3 hours and 11 minutes, producing an RMS of R peak displacements of 49.051 ms for a period from 30 s to 23 minutes (SDNN of real RRI was 88.263) as the most optimal solution. For a quinquagenarian male, 17533 evaluations took about 4 hours and 43 minutes, resulting in an optimal solution with RMS of R peak displacements of 27.925 ms (SDNN of real RRI was 32.038). None of the most optimal solutions had a penalty score. **Table 1** discloses the most optimal sets of personalized parameters.

The main result of our proposed concept is the time course of the R peak displacements, shown in **Figure 12** and **Figure 13** parts C and F for each subject. The R peak displacements of the tricenarian male had a relatively stable baseline despite the real HR baseline drifts (**Figure 12A**), even during shifts between resting and slow breathing periods. The R peak displacements had pulsations of about 0.1 Hz at multiple time intervals (**Figure 12F**) that often resemble ABP (**Figure 12E**) with a time delay of about 2.2 s. Furthermore, some R peak displacements overshot the mentioned pulsations just for a single RRI, and these momentary heart decelerations were up to 280 ms compared to the expected by the personalized HR model (marked with red arrows in **Figure 12F**, e.g., the largest one appeared at 82 s). These decelerations often occurred at the rising part of slow R peak displacement pulsations after the previous drop of ABP with SNA burst (**Figure 12E**) and expiration (**Figure 12D**). In fact, in both subjects, SNA is generated more often and intensely at drops in ABP via withdrawal of the modeled baroafferent activity (**Figure 12E**). Baseline drifts existed in R peak displacements of the quinquagenarian male (**Figure 13C**). Here, R peak displacements did not resemble ABP in general. However, a few morphological similarities are visible, e.g., about 0.1 Hz pulsations at about 970–990 s and baseline drop at about 1060–1090 s intervals (compare **Figure 13** parts E and F). In both cases, the time course of the R peak displacements differed from respiration, especially in the quinquagenarian male. The fast virtual HR variations mimicked respiration and real HR patterns (**Figure 12D** and **Figure 13D**). Therefore, three main components of the R peak displacements may be visible to different individuals even during the same resting condition: very slow baseline drifts (<0.1 Hz), about 0.1 Hz pulsations, and momentary fast irregularities.

**Figure 12.**
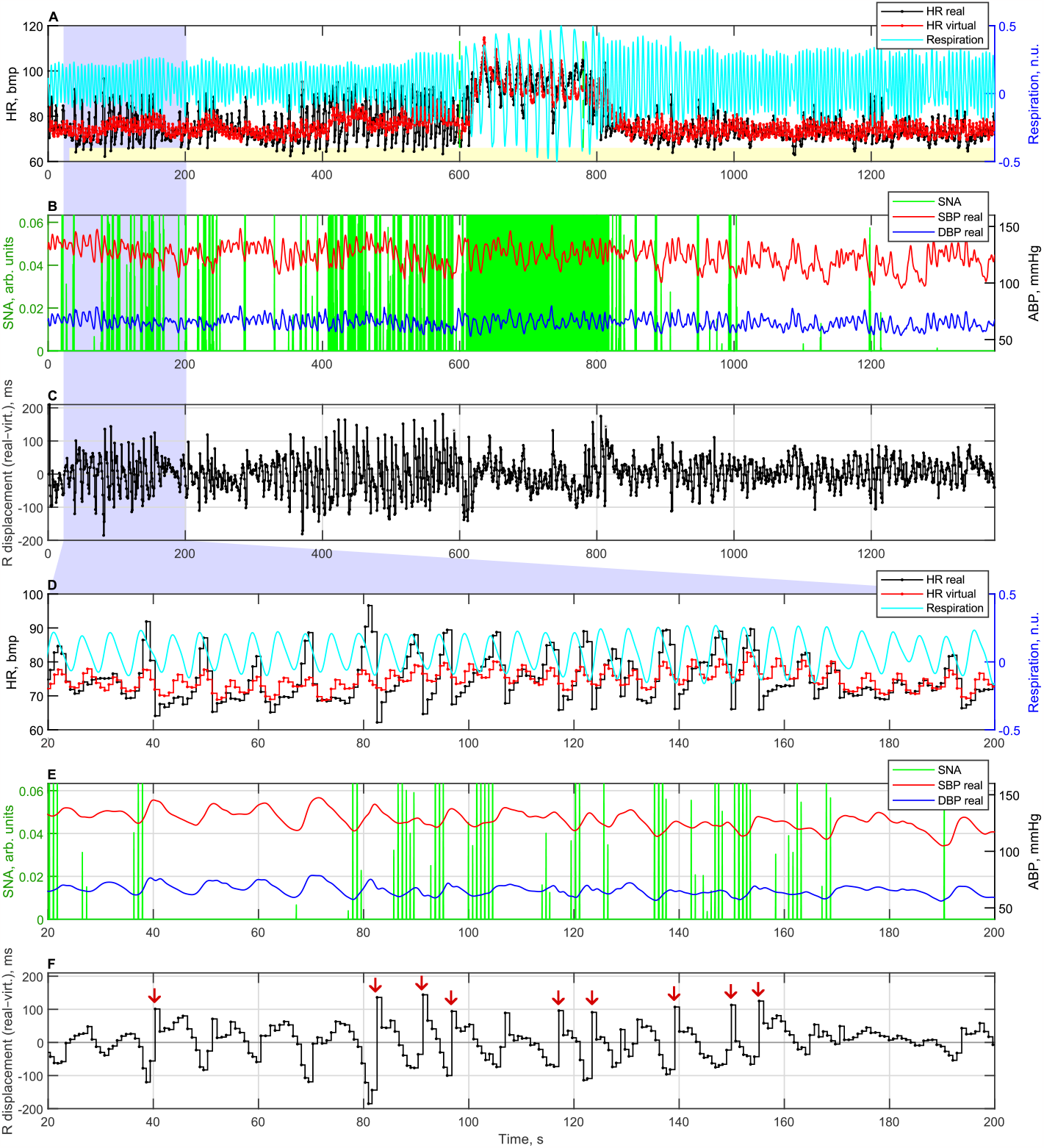
Psychophysiological data of a tricenarian male: time course of the real and virtual heart rate (HR), respiration (A and D parts), the modeled sympathetic nerve activity (SNA), and the processed real arterial blood pressure (ABP) (B and E parts), R peak displacements (C and F parts). The A, B, and C parts are enlarged in D, E, and F. The period of slow respiration condition was from 600 to 780 seconds (green dashed marks in A part); other periods are in the resting state. HR model calibrated for a period from 30 to 1380 seconds (light yellow strip at the bottom of A part). Other abbreviations: DBP, diastolic blood pressure; SBP, systolic blood pressure.

**Figure 13.**
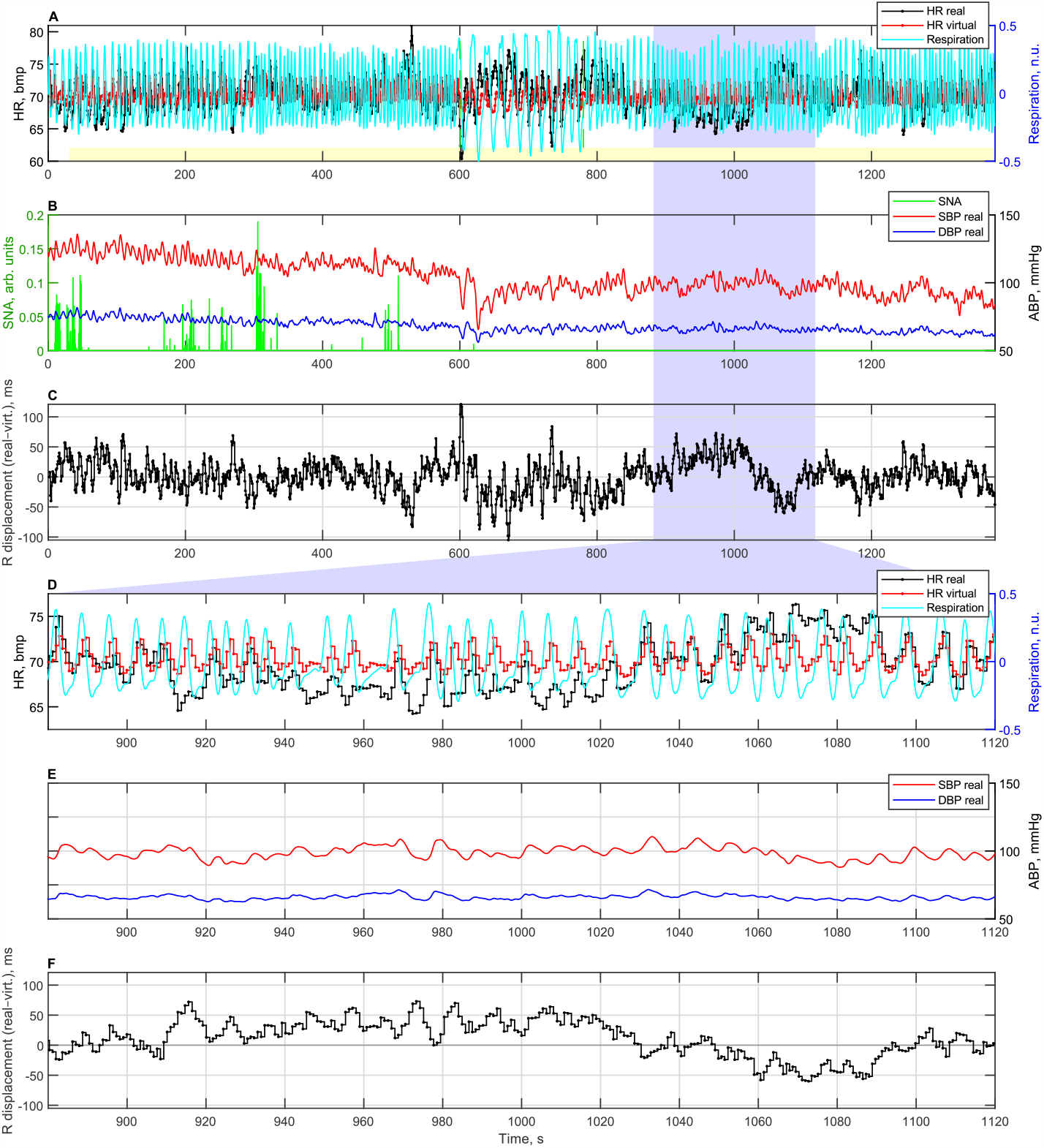
Psychophysiological data of a quinquagenarian male: time course of the real and virtual heart rate (HR), respiration (A and D parts), the modeled sympathetic nerve activity (SNA), and the processed real arterial blood pressure (ABP) (B and E parts), R peak displacements (C and F parts). The A, B, and C parts are enlarged in D, E, and F. The period of slow respiration condition was from 600 to 780 seconds (green dashed marks in A part); other periods are in the resting state. HR model calibrated for a period from 30 to 1380 seconds (light yellow strip at bottom of A part). Other abbreviations: DBP, diastolic blood pressure; SBP, systolic blood pressure.

## 4 Discussion

This article describes a concept of computerized physiology modeling that integrates physiology knowledge and cutting-edge computing technologies into virtual models. Virtual models allow not only a more efficient investigation of physiological mechanisms, predict multiple influences, but also enable the application of the modeling results to biosignal and data processing, research, and clinical decision-making (Pruett et al., 2020). This is especially relevant for the progress of personalized medicine, prediction, and prevention. Modeling of HR regulation is an essential part of such approaches as virtual physiological human (www.vph-institute.org) or virtual patients for *in silico* trials (Sinisi et al., 2021). Article contributes to two main research problems in modeling HR regulation: 1) how to integrate the abundant knowledge of physiology, psychology, and biophysics in the form of formal models that are easily accessible to the user of computerized systems; and 2) how to enrich these models with additional empirical personal data, thereby refining the models and personalizing the decisions they could support.

The specific purpose of this paper was to present the concept of a computational method that could identify personalizable HR regulation parameters and capture the beat-by-beat variations (R peak displacements) that are not expectedly attributable to lower order HR regulation mechanisms – RSA and cardiac baroreflex. To our knowledge, such a method incorporating a multilevel model of HR regulation, designed to be fitted to empirical data and seeking to extract residual HR irregularities, has not yet been proposed. A valuable feature of the model is the presentation of neurophysiological regulatory mechanisms in the visual form of SIMULINK block diagrams, which makes the model and simulation easily interpretable.

Most of the models available thus far only allow generalized modeling of humans without the possibility of tailoring the model to a specific person. Some attempts to create patient-specific models (Gray and Pathmanathan, 2018) have been proposed, however the trend is to create more universal models suitable for either normal or pathological (i.e., personalized) conditions, paving the path from the virtual physiological human to the clinic (Hoekstra et al., 2018). A popular PNEUMA model (Hsing-Hua Fan and Khoo, 2002; Cheng et al., 2010) is highly elaborated and can simulate multiple influences even within the heart period, however, this model is too complex and would be too slow to search for parameter combinations for beat-to-beat personalization. Some newer models, relying on the knowledge of lower-level mechanisms, propose more elaborated neural regulations at relatively higher levels. For instance, the Park et al. (2020) model included a more detailed parasympathetic activity regulation via NTS, NAmb, and dorsal motor nucleus of the vagus, nevertheless, this is still about regulation at the lower-order medulla oblongata level. Although HR regulation models involving multiple higher brain structures have already been described in the literature (Thayer and Lane, 2009; e.g., Smith et al., 2017), they lack a time axis and mathematical formulations.

To evaluate HR regulation influences, particularly those attributable to higher-order regulations, a part of a model related to lower-order regulations must be adequate to the *in vivo* HRV. In particular, our proposed model could simulate RSA and cardiac baroreflex, mimicking the morphology of the real HR. Our model personalization system found sets of parameters of the HR model suitable for two psychophysiological conditions – resting and slow breathing – for two males. Both sets were useful for generating R peak displacements that are different from the respiration morphology, suggesting that the architecture of the RSA mechanism is adequate in our HR model.

Our proposed concept reveals the possibility of separating three components of R peak displacements – tonic, spontaneous, and 0.1 Hz changes – and analyzing them separately. The *tonic component*, like the baseline drifts seen in the quinquagenarian male, could be associated with slow changes in the mental, physiological, and humoral state. The *spontaneous changes* in the R peak displacements seen in the tricenarian male recording, while preliminary, suggest other factors here, they can also indicate sudden sporadic changes in parasympathetic activity from higher-order brain areas, possibly partly due to additional interaction between baroinformation and RSA. The *0*.*1 Hz component* may be related to Mayer waves or insufficient model adequacy in simulating the baroreflex from ABP. There is a 2.2 s delay of R displacements relative to the ABP (see **Figure 12 E** and F) that cannot be fitted to faster vagal or slower sympathetic baroreflex mechanisms, therefore adding new mechanisms and their interactions or implementing additional inputs could improve the existing model. Nevertheless, interesting insights can come from analyzing components of R peak displacements extracted by our concept, even without modifications.

Our proposed model-based computational method has limitations and simplifications that may be improved. One limitation is assuming a constant tonic level of ANS activity as higher-order neural control. However, an effective input from the higher-order brain is expected at about 400–600 ms after the R peak, as suggested by our previous study (Baranauskas et al., 2021). Another is that our model currently does not simulate orthostatic hypotension and is intended to be used when the person remains in the same posture. Furthermore, unexplained HR variations are provisionally attributed to higher-order brain regulation in our proposed approach, even though they could be attributed to any heart regulation level. The results described do not claim to validate the model – they were just for illustrative purposes to prove that the concept just works. Validation of the model could be performed using larger samples of physiological recordings, preferably compared with other methods and evaluated statistically.

Nevertheless, our proposed approach could help investigate brain-heart interactions by enhancing the ratio between influences from the brain and other HR regulation mechanisms. Future studies could try to discriminate sources of displacements of R peaks, such as higher-order neural or humoral, either sympathetic or parasympathetic. The model-based computational system should also be tested on a broader sample of subjects to reflect sex, age, and specific illnesses; for example, it is known that cardiac vagal baroreflex is relatively more pronounced in females and sympathetic baroreflex in males (Kim et al., 2011).

The model-based concept could be further applied to the *in-silico* analysis of heart regulation in healthcare, medical diagnostics, psychology, and research on heart-brain interactions. The model parameters themselves could be valuable in assessing RSA and the baroreflex. R displacements could help control RSA and baroreflex effects in stress, fatigue, emotion studies, or biofeedback systems while considering tonic, phasic, or spontaneous changes in HR. The model could also help develop diagnostic and screening biomarkers (e.g., sympathovagal balance) based on HRV data retrieved from wearable devices. The developed computational system that integrates the psychophysiological mechanisms of HR regulation in a formalized personalized model may facilitate further research and reveal new features that would be useful in practice.

## Acknowledgments

The authors thank all the volunteers for their participation in the study. The authors thank the Institute of Biomedical Electronics of the Vienna University of Technology for laboratory equipment for psychophysiological recordings. We thank Lena Kummer for her assistance in the experimental study. We thank Ana Rita Alves dos Santos Rodrigues for English language reviewing.

## Conflict of Interest

The authors declare that the research was conducted in the absence of any commercial or financial relationships that could be construed as a potential conflict of interest.

## Author Contributions

MB and RS contributed to conception. AL and MB contributed to funding acquisition. EK designed the pilot study for data collection. EK and MB did pilot study and collected the data. MB cared about modeling, software, data analysis, interpretation, visualization of results, and wrote the first draft of the manuscript. All authors contributed to manuscript revision, read, and approved the submitted version.

## Ethics statement

Ethical approval was not provided for this study on human participants because pilot recordings are used as the basis for the ethic application (namely, for the estimation of the effect size and thus for the necessary size of the clinical study). The participants provided their written informed consent to participate in this study. Written informed consent was obtained from the individuals for the publication of any potentially identifiable data included in this article.

## Funding information

This work was funded by the European Social Fund under No 09.3.3-LMT-K-712 “Development of Competences of Scientists, other Researchers and Students through Practical Research Activities” measure.

## Data Availability Statement

The raw data supporting the conclusions of this article will be made available by the authors, without undue reservation.

## Notes

https://doi.org/10.5281/zenodo.7765459

### Competing Interest Statement

The authors have declared no competing interest.

### Summary of Updates

English language revised and improved.

